# Effects of trap confinement on personality measurements in two terrestrial rodents

**DOI:** 10.1101/723403

**Authors:** Allison M. Brehm, Sara Tironi, Alessio Mortelliti

## Abstract

In recent years individual differences in the behavior of animals, or personalities, have been shown to influence the response of individuals to changing environments and have important ecological implications. As researchers strive to understand and predict the responses of individuals and populations to anthropogenic changes, personality studies in wild populations will likely continue to increase. Studies of personality in wild populations often require that animals are live-trapped before behavioral observation can occur; however, it is unknown what impact live trapping may have on the behavior of trapped individuals. Specifically, if the duration of trap confinement directly influences behavior, then by obtaining wild animals through live-trapping are we confounding the very measurements we are most interested in? To investigate this question, we performed a study using two small mammal species. We positioned high-definition trail cameras on Longworth small mammal traps in the field to observe capture events and record the time of capture. We then measured personality in captured deer mice (*Peromyscus maniculatus*) and southern red-backed voles (*Myodes gapperi*) using three standardized tests. With a repeatability analysis, we confirmed which behaviors could be considered personality traits, and through linear and generalized linear models, we found that the time an animal had spent confined to a trap before testing did not affect the majority of behaviors exhibited. Our results showed two weak behavioral effects of confinement duration on boldness and docility depending on whether an individual had been trapped previously. Our results suggest that personality measurements of wild, trapped small mammals are not determined by trapping procedures, but that researchers should control for whether an animal is naïve to trapping during analysis.

## Introduction

Over the past few decades, the acknowledgement that many species of animals display consistent individual differences in behavior, or *personalities*, has become widespread (1–4). Personalities are heritable (5), have consequences for fitness (6–9), and can limit the ability of individuals to exhibit behavioral plasticity (10) resulting in trade-offs where certain personality types perform well in some ecological contexts but not in others (11). Because individual personalities can determine the response of individuals to changing environments (12,13) and have important ecological implications (14–16), personality studies in wild populations will likely continue to increase as researchers strive to understand and predict the responses of individuals and populations to anthropogenic changes (17–20).

Studies of personality in wild populations usually require that wild animals are live-trapped so that one or more standardized behavioral tests can be undertaken (21–24) but see (25) for a method of personality observation in non-captured animals. Because being trapped may induce stress (26–31), the process of capturing animals and subsequently measuring their personality offers additional challenges. Specifically, the stress of being trapped might influence the behaviors exhibited by wild animals, confounding the very phenomena we are investigating.

Several studies have explored the relationship between live trapping and the stress response of animals (29–31), and it is generally accepted that the stress of being captured releases glucocorticoids into the bloodstream (32). Glucocorticoids act to elevate breathing rate, heart rate, and blood pressure (29) which, following exposure to the threat of a predator attack, stimulates the mobilization of energy to facilitate an escape. When an animal is confined to a trap, however, this prolonged stressor may result in higher concentrations of glucocorticoids after longer durations spent in a trap (30), perhaps impacting behaviors exhibited during routine behavioral tests such as grooming, time spent moving, etc. (33–35). Thus far, studies looking to assess this phenomenon have focused on the hormonal/physiological response to trap-induced stress and results have been mixed (29,31,36). For example, live trapping does induce an initial stress response in southern red-backed voles (*Myodes gapperi*) and meadow voles (*Microtus pennsylvanicus*), but longer times spent in traps do not correlate with increased stress levels (29,36). In contrast, studies found that in deer mice (*Peromyscus maniculatus*) and North American red squirrels (*Tamiasciurus hudsonicus*) prolonged time spent in traps was positively correlated with stress hormone levels (31,36). In either scenario, it is unknown whether the time spent in traps may produce a behavioral response, since a change in stress hormones doesn’t necessarily precede a change in behavior. If confinement duration affected the behavior exhibited during routine testing, this would require studies using personality data from trapped animals to control for confinement duration. This could be done by: checking traps more frequently, recording the time of capture (obtained using videos from camera traps placed on live traps) then controlling for the duration using imposed covariates in analysis, or using devices that signal when a capture has been made so that animals can be removed promptly (37,38). Empirical evidence is needed to explore the relationship between the time spent in a trap and behavioral response.

The objective of this study was to assess whether personality measurements obtained from live-trapped individuals are being confounded by the amount of time spent inside of a trap. Specifically, we sought to determine whether confinement duration affects the behaviors exhibited in routine behavioral tests. To meet this objective, we conducted a field experiment focused on the deer mouse (*Peromyscus maniculatus*) and the southern red-backed vole (*Myodes gapperi*), which have been the subject of previous personality studies (16,39). Using high-definition trail cameras positioned on Longworth small mammal traps in the field, we quantified the duration of time that individuals had spent inside a trap before behavior was observed in standardized behavioral tests the following morning. We explored these data to see whether behaviors exhibited in behavioral tests varied with the time spent inside the trap.

Results from this study will have implications for researchers who measure personality following the live-capture of an animal. These results will highlight whether we should take additional steps to ensure that our behavioral measurements are accurate and not unduly influenced by the trapping.

## Materials and methods

### Study site and small mammal trapping

This study was conducted in the Penobscot Experimental Forest (PEF, 44 51’ N, 68 37’ W) at the southern edge of the Acadian forest in east-central Maine, USA. This experimental forest consists of forest units chosen at random and logged separately with varying silvicultural treatments (minimum of two replicates per treatment). Management units average 8.5 ha in area (range 8.1–16.2 ha) and nearly 25 ha of forest (retained in two separate units) serves as reference and has remained unmanaged since the late 1800s (39,40).

We implemented a large-scale mark-recapture study on six trapping grids (Figure 1): two control (located in reference forest) and four experimental (two replicates in even-aged forest units and two in units treated with a two-stage shelterwood with retention). Trapping grids were 0.81 ha in area and consisted of 100 flagged points spaced 10 m apart. One Longworth trap was positioned at each flagged point. Traps were bedded with cotton and baited with a mixture of sunflower seeds, oats, and freeze-dried mealworms. We positioned trapping grids close to the center of the management unit to minimize edge-effects (mean distance between grids was 1.44 km; greater than the movements of our study species). We trapped at each trapping grid for three consecutive days and nights and checked traps each morning and evening. Trapping occurred once per month for five consecutive months each year (June–October 2016, 2017, 2018).

**Figure 1.**
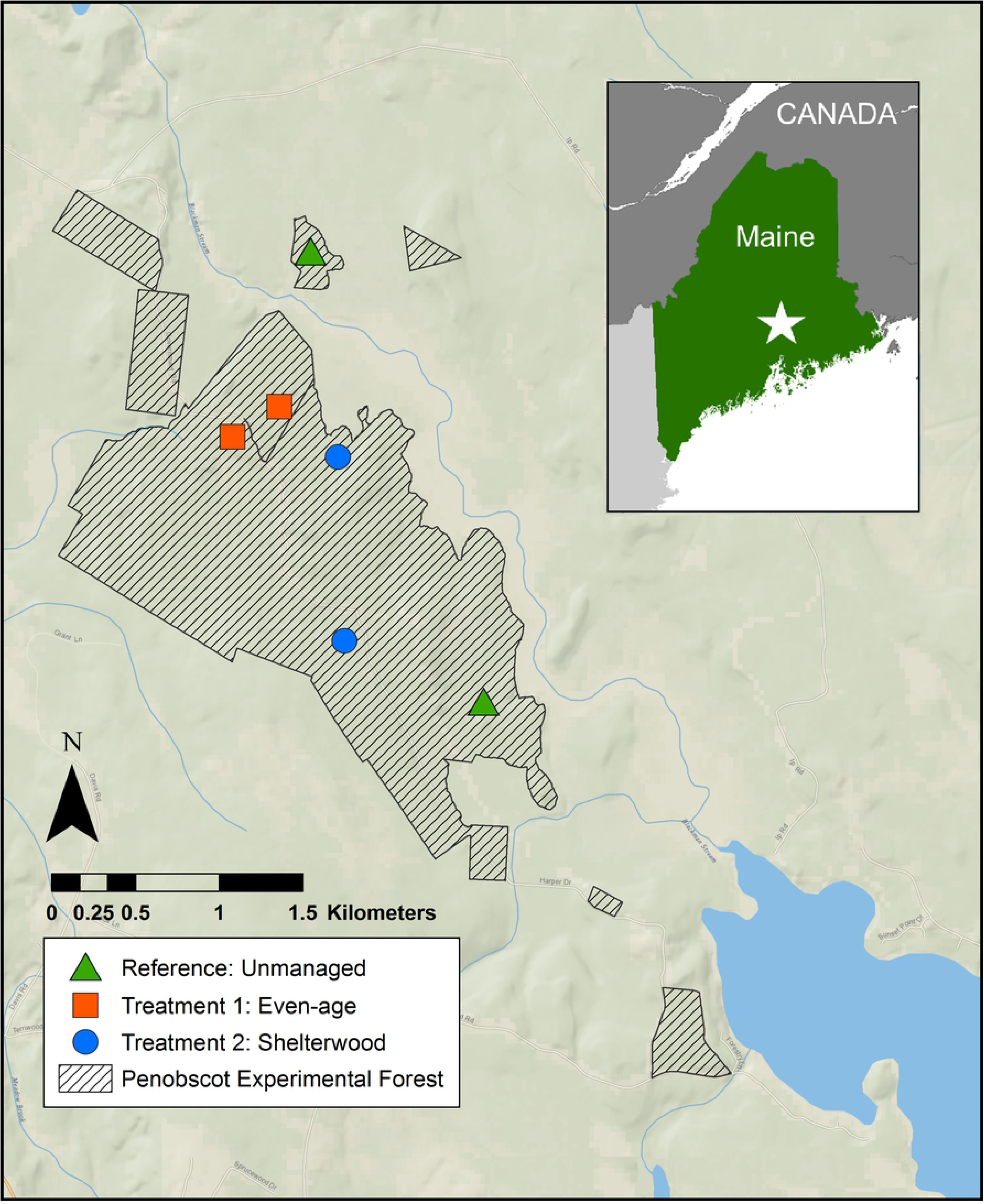
Map of our study area at the Penobscot Experimental Forest, Maine U.S.A.

### Behavioral tests

We used three standard behavioral tests to measure personality of trapped individuals (Figure 2): an *emergence* test to assess boldness (33,41), an *open-field* test to measure activity and exploration in a novel environment (42,43), and a *handling bag* test to measure docility and the response to handling by an observer (23,44–46). Behavioral tests were performed in the order above prior to handling or marking. All tests and processing occurred at a base area in the home grid of the focal individual.

**Figure 2.**
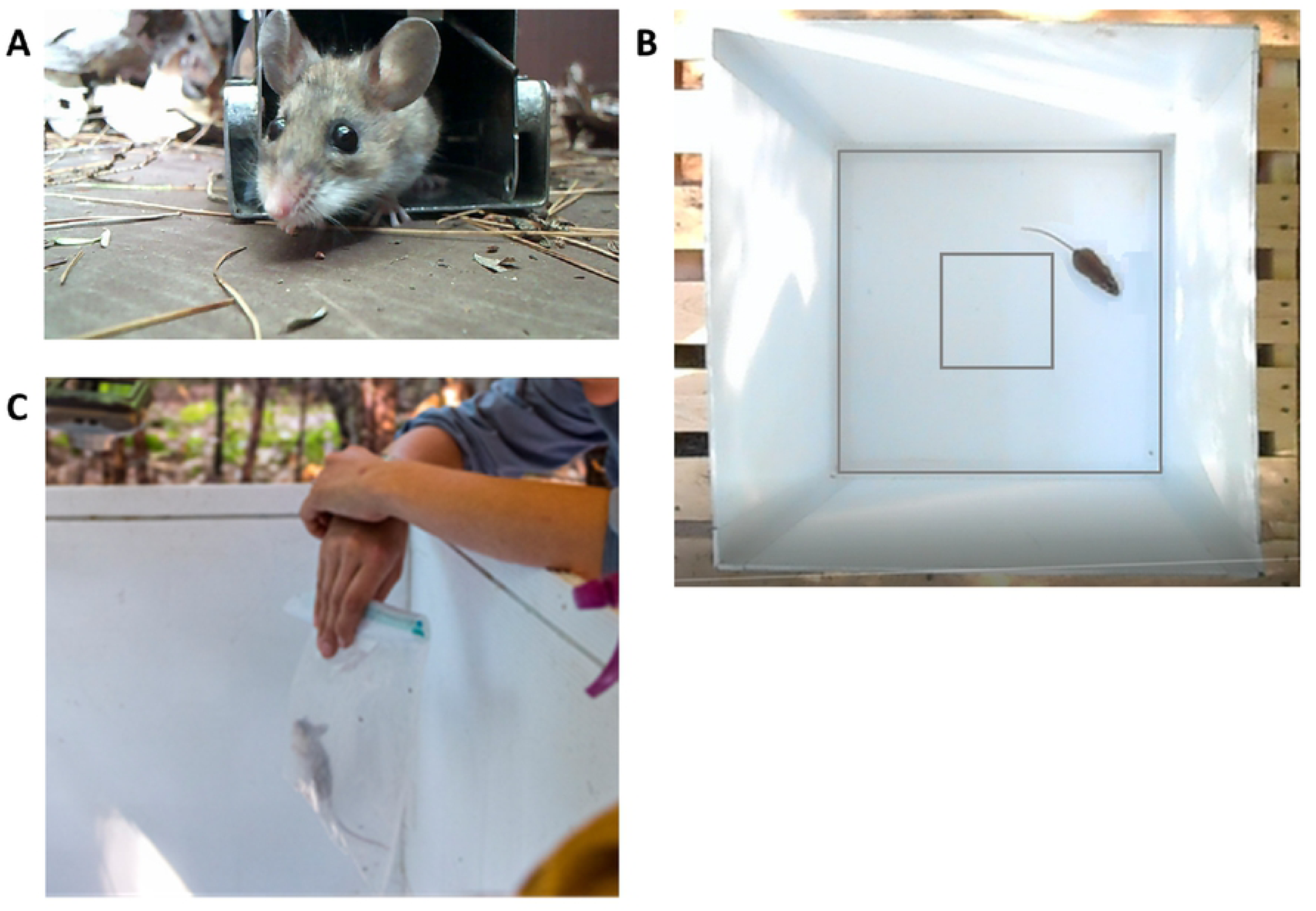
Three behavioral tests used to assess personality of deer mice (*Peromyscus maniculatus*) and southern red-backed voles (*Myodes gapperi*). (A) An individual emerges from a Longworth trap in an emergence test. (B) An individual in motion during an open-field test. (C) An observer suspends an individual over a controlled arena during the handling bag test.

Behavioral tests were performed as follows: first, the animal was transferred directly from the trap of capture into a clean, empty Longworth trap. This trap was then placed on the floor of a box sized 46 x 46 x 60 cm (placed underneath a tarp to control for light levels and perceived canopy cover). To create a more natural environment, the inside of the box had been painted a light brown with a small amount of debris (dead leaves and pine needles) placed on the floor. A digital camera (Nikon CoolPix S3700) was mounted facing the opening of the Longworth trap, and the observer locked the trap door open before leaving the test area. After three minutes, the observer returned and ended the test. Individuals were caught in a plastic bag and then released into the center of the open-field arena. A five minute open-field test was performed in an arena (46 x 46 x 50 cm), placed on a level platform with perceived canopy cover controlled (39), and a mounted digital camera (Nikon CoolPix S3700) recorded the test. After five minutes, an observer ended the recording, caught the animal in a plastic bag, and performed a handling bag test by suspending the bag into the open-field arena to control the visual surroundings. The observer measured the proportion of time that the individual spent immobile during one minute (referred to as handling time hereafter). Traps used for emergence tests and the open-field test box were cleaned thoroughly with 70% isopropyl alcohol and wiped with a dry cloth in-between all tests. Behavioral tests were performed once monthly to ensure that animals would not habituate to the tests.

After the completion of the behavioral tests, we anesthetized animals with isoflurane and inserted PIT tags (Biomark MiniHPT8) subcutaneously at the midback. Animals were also marked with a small animal ear tag (National Band, Style 1005-1) and a distinctive haircut. We recorded sex, body mass (measured using a 100 g Pesola Lightline spring scale), body length, tail length, reproductive status, and age class (juvenile, subadult, or adult). Animals were released at the exact site of capture post-processing.

To quantify behavior from videotaped emergence and open-field tests, recordings were played back in the laboratory. For emergence tests, an observer recorded the following: whether or not the animal emerged (defined as all four feet having left the trap), the latency to emerge, and the total time spent at the end of the tunnel before emerging. Open-field tests were analyzed using the behavioral tracking software ANY-maze © (version 5.1; Stoelting CO, USA). For the remainder of analyses, we focused on a reduced number of non-redundant behavioral variables (16). See Table 1 for a complete list of the behaviors used.

**Table 1.**
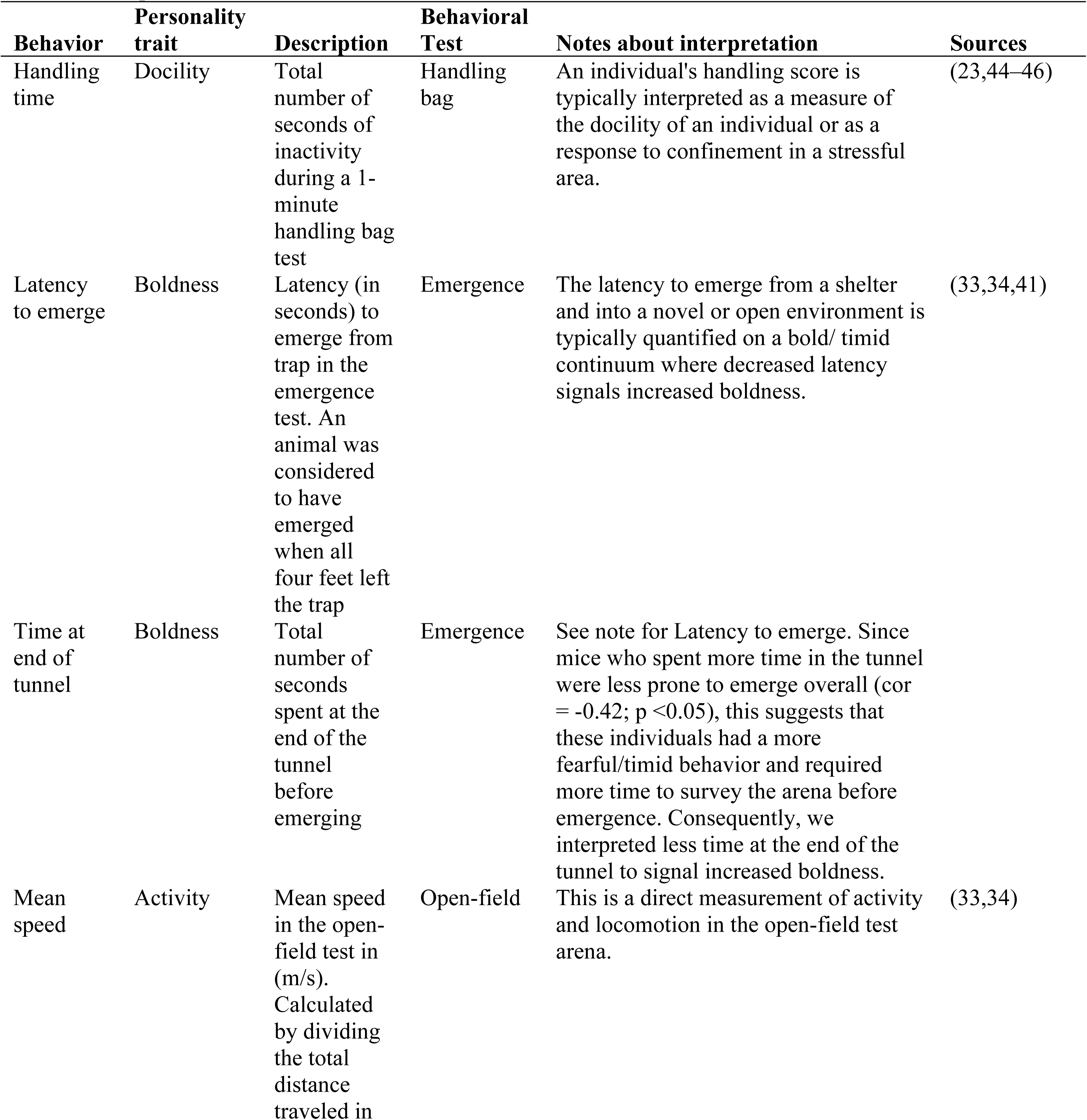
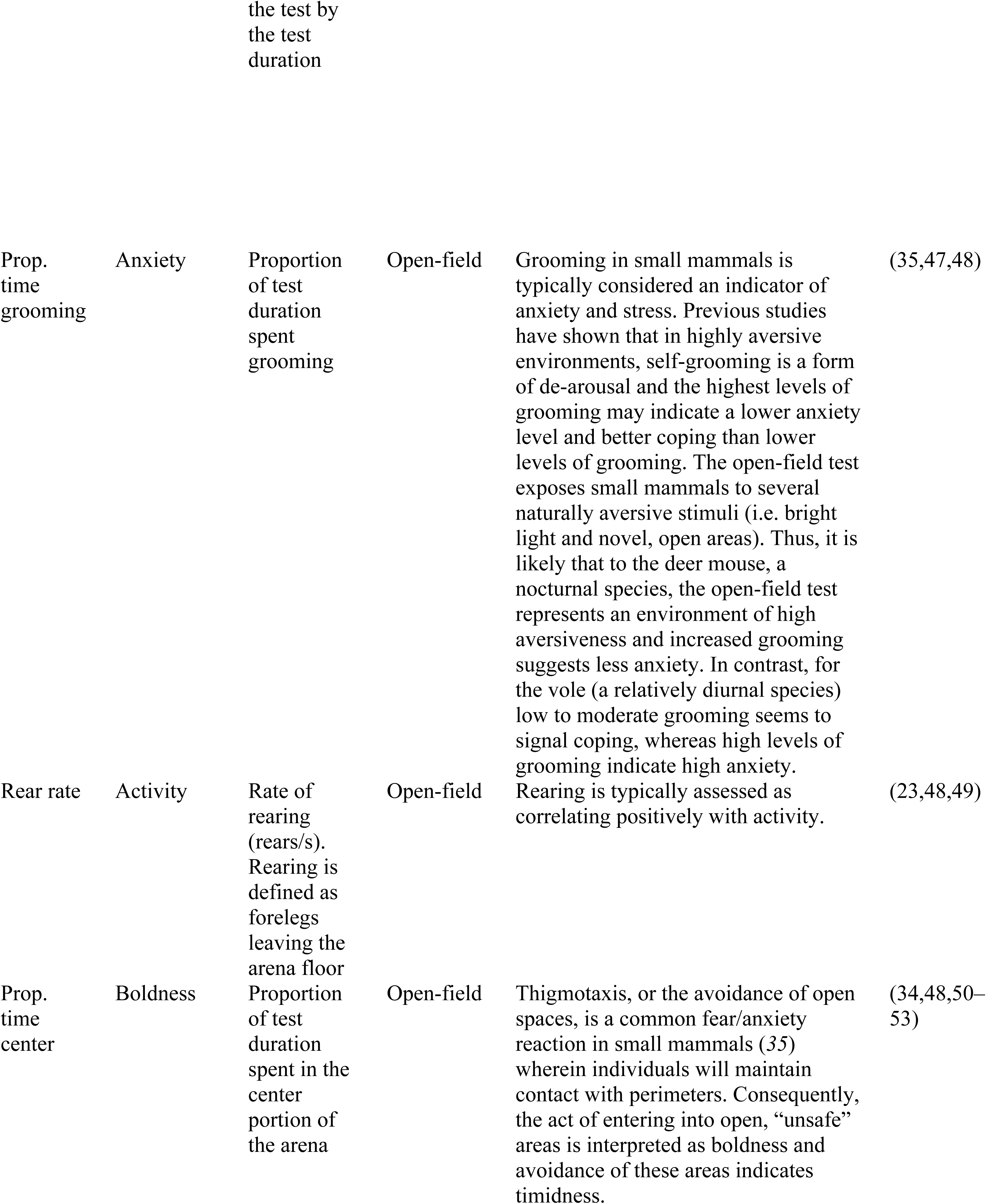
Personality variables measured in the deer mouse (*Peromyscus maniculatus*) and the southern red-backed vole (*Myodes gapperi*). Provided are: the behavior, description, behavioral test it was measured using, notes on interpretation, and a non-exhaustive list of references.

### Monitoring capture events

To observe the event of an individual’s capture and calculate the time spent inside the trap before behavioral testing, we positioned camera traps (Bushnell NatureView HD 119740) facing the door of the Longworth trap and its surroundings. Cameras were positioned ∼50–100 cm from the trap at a height of ∼50 cm. 13 camera traps were used in total and were positioned on a subset of the 100 available trap locations (Figure 3). We chose camera locations to optimize the chance of observing capture events (hence, we chose trap locations that had successful captures during the previous month’s trapping session). Cameras were positioned simultaneously with Longworth traps and were kept active for the same duration as the traps (three consecutive days and nights at each study grid). We monitored Longworth capture events using camera traps from July–October 2018 (936 total camera trap nights). Cameras were programmed to record a one-minute video whenever movement was perceived (with a one second delay between videos). Because camera traps occasionally fail to detect movement, we also programmed them to take a one-minute video once per hour (the “field scan” setting). This allowed us to approximate the hour of capture in an instance where the camera failed to trigger at the capture event.

**Figure 3.**
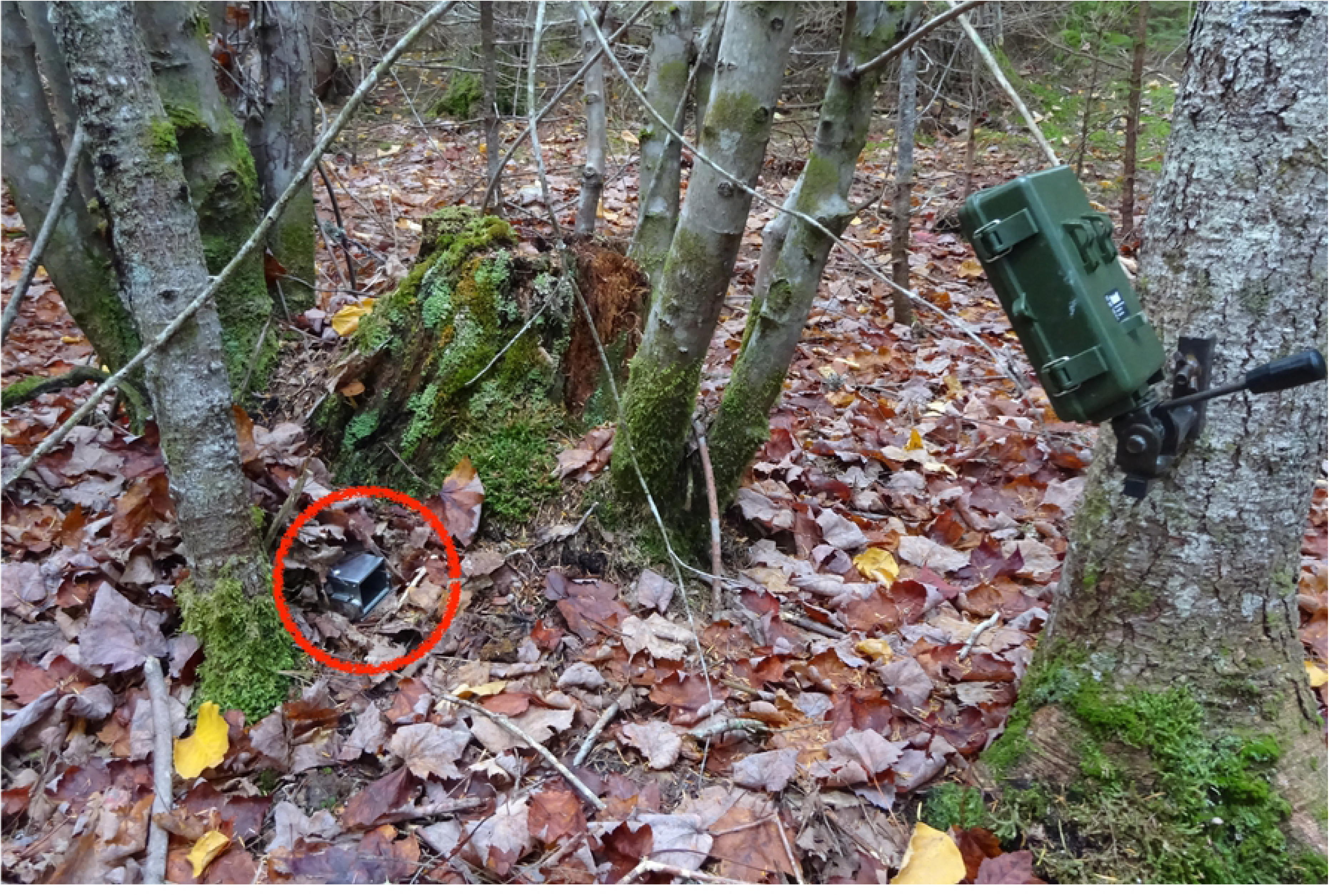
A camera trap (Bushnell NatureView HD) monitors a Longworth trap in the field.

Videos of capture events were played back in the laboratory, and an observer identified the individual by pairing the information of the date and trap with available capture data. The observer then recorded the time that the individual entered the trap and calculated the total time (in minutes) spent inside the trap before behavioral testing (taken from the time stamp of the open-field video for consistency). This variable will be referred to hereafter as “time in trap”. See S1 Video and S2 Video in the supporting information for examples of observed capture events.

**S1 Video. Observed capture event of a southern red-backed vole (*Myodes gapperi*).**

**S2 Video. Observed capture event of a deermouse (*Peromyscus maniculatus*).**

### Data analysis

To determine which behaviors could be considered personality, we first performed a repeatability analysis on the behavioral variables obtained from the emergence, open-field, and handling bag tests (54,55). For this analysis, we used data from our study population collected during the 2016, 2017, and 2018 field seasons. We used R version 3.4.1 (56) and package *lme4* (57) to run mixed-effects models and included potential confounding factors as covariates in the models. Specifically, we included sex, body condition (calculated using the scaled mass index (58)), silvicultural treatment, trapping session (June–October), and trapping year (2016, 2017, or 2018). Individual identity was included as a random intercept in the models to account for the proportion of the variance that can be attributed to differences among individuals (59). As response variables, we used the behavior of interest and ran separate mixed-effects models for each behavior of interest. We assessed normality by visually inspecting Q–Q plots and histograms of the residuals, and by plotting the fitted values against the residual values (60). We logit-transformed the response variable when it was a proportion (59,61) to meet the assumption of normality. We then calculated the adjusted repeatabilities and associated confidence intervals (55,62–64) using methods described in detail by (16,39).

Once it was determined which behaviors were repeatable and could, therefore, be considered personality, we tested the hypothesis that these behaviors would be influenced by the time spent inside the Longworth trap before behavioral testing. We used a nested hypothesis testing approach (65) using linear models and generalized linear models with the repeatable behaviors as response variables. In the instances where we had repeated measures from the same individual (because we caught their capture on a camera trap in subsequent trapping sessions), we used only the first event (18 out of 92 individuals). Again, proportional response variables were logit-transformed to meet the assumptions of normality, and count variables were examined using generalized linear models with a poisson or negative binomial family (depending on dispersion).

We introduced predictor variables one by one to build a base model to control for most of the variability in the data. Predictor variables included sex, body condition, silvicultural treatment, trapping session, body mass, and a variable termed “naïve” which controlled for whether the animal had been captured previously or was naïve to trapping. Models containing each of these variables alone were compared to the null model using the Akaike information criterion corrected for small sample size (AICc) (65,66) and models within 2.0 ΔAICc of the top model were considered to have equal support. If more than one variable was better than the null, a model including multiple additive effects was explored. Once this base model was built, we compared this model to the same model with the addition of the variable “time in trap” to see whether this addition improved the model by AICc. Previous research has shown that males and females may respond differently to trap-induced stress (31), so we subsequently tested for an interaction between the time spent in the trap and sex. Last, to test the hypothesis that individuals who are naïve to trapping may be impacted by the time spent inside the trap differently than individuals who have been captured previously, we tested for an interaction between time spent in the trap and the variable “naïve”.

### Ethical note

Animal trapping, handling, and marking procedures were approved by the University of Maine’s Institutional Animal Care and Use Committee (IACUC number A2015_11_02). Animals were anaesthetized with isoflurane prior to tagging, and tagging equipment was sanitized with 70% isopropyl alcohol in between animals. All small mammal handling was performed by trained researchers, and all efforts were made to minimize suffering by small mammals.

## Results

We examined behavioral data from standardized tests for 1791 observations from 603 individual deer mice and 1558 observations from 529 individual red-backed voles, and we found all behavioral variables to be significantly repeatable, with a mean repeatability value of 0.81 for deer mice and 0.78 for voles (Table 2). This indicates that these behaviors can be considered personality (55,67). The mean 95% confidence intervals for these values were (0.79, 0.84) and (0.74, 0.81), respectively (Table 2). The number of observations and individuals shown in Table 2 differ for behavioral variables obtained from the emergence and handling bag tests since these tests were not performed in 2016. The mean number of repeated observations per individual was approximately three for both deer mice and red-backed voles.

**Table 2.** Repeatability estimates for target behaviors measured in three behavioral tests (handling bag, emergence, and open-field) in deer mice (*Peromyscus maniculatus*) and southern red-backed voles (*Myodes gapperi*).

| Behavioral Variable | Mean | Range | Repeatability (95% CI) | Observations | Individuals |
| --- | --- | --- | --- | --- | --- |
| ***P. maniculatus*** |  |  |  |  |  |
| Handling time | 15.41 | (0, 60) | **0.836 (0.807, 0.862)** | 1122 | 376 |
| Latency to emerge | 27.17 | (0, 180) | **0.812 (0.780, 0.842)** | 1122 | 376 |
| Time at end of tunnel | 7.15 | (0, 180) | **0.863 (0.841, 0.884)** | 1122 | 376 |
| Mean speed (m/sec) | 0.10 | (0, 0.25) | **0.832 (0.809, 0.853)** | 1791 | 603 |
| Prop. time grooming | 0.11 | (0, 0.96) | **0.762 (0.735, 0.792)** | 1791 | 603 |
| Rear rate | 0.19 | (0, 0.68) | **0.809 (0.785, 0.831)** | 1791 | 603 |
| Prop. time center | 0.03 | (0, 0.73) | **0.775 (0.747, 0.804)** | 1791 | 603 |
| ***M. gapperi*** |  |  |  |  |  |
| Handling time | 47.77 | (0, 60) | **0.675 (0.62, 0.726)** | 940 | 305 |
| Latency to emerge | 34.94 | (0, 180) | **0.831 (0.799, 0.859)** | 940 | 305 |
| Time at end of tunnel | 12.05 | (0, 180) | **0.823 (0.791, 0.851)** | 940 | 305 |
| Mean speed (m/sec) | 0.05 | (0, 0.20) | **0.792 (0.765, 0.818)** | 1558 | 529 |
| Prop. time grooming | 0.06 | (0, 0.81) | **0.729 (0.694, 0.764)** | 1558 | 529 |
| Rear rate | 0.09 | (0, 0.56) | **0.770 (0.739, 0.801)** | 1558 | 529 |
| Prop. time center | 0.04 | (0, 0.99) | **0.827 (0.805, 0.850)** | 1558 | 529 |

Repeatability was calculated from univariate mixed-effect models with identity included as a random effect. Parametric bootstrapping was used to calculate 95% confidence intervals. See Methods for more information. Significant repeatability estimates are shown in bold.

In the majority of models (∼86%) predicting behaviors exhibited in standardized tests, the top model did not include “time in trap”. Instead, out of the predictor variables considered (sex, body condition, silvicultural treatment, trapping session, body mass, and a variable termed “naïve” which controlled for whether the animal had been captured previously or was naïve to trapping) behaviors in deer mice were predicted by trapping session and body mass (Table 3, Figure 4a-b). Deer mice with greater body mass showed longer latencies to emerge from the emergence test and the proportion of time spent grooming in the open-field test correlated positively with trapping session (β = 0.26, SE = 0.08, rsq = 0.20 and β = 0.58, SE = 0.16, rsq = 0.23, respectively). In two cases, (once for deer mice and once for voles) the top model included an interaction between “time in trap” and whether or not the individual was naïve to trapping (Figure 4c-d). Model fit was relatively low for top models (excluding those where the top model included only an intercept), with an average multiple R-squared value of 0.23 (Table 3).

**Table 3.**
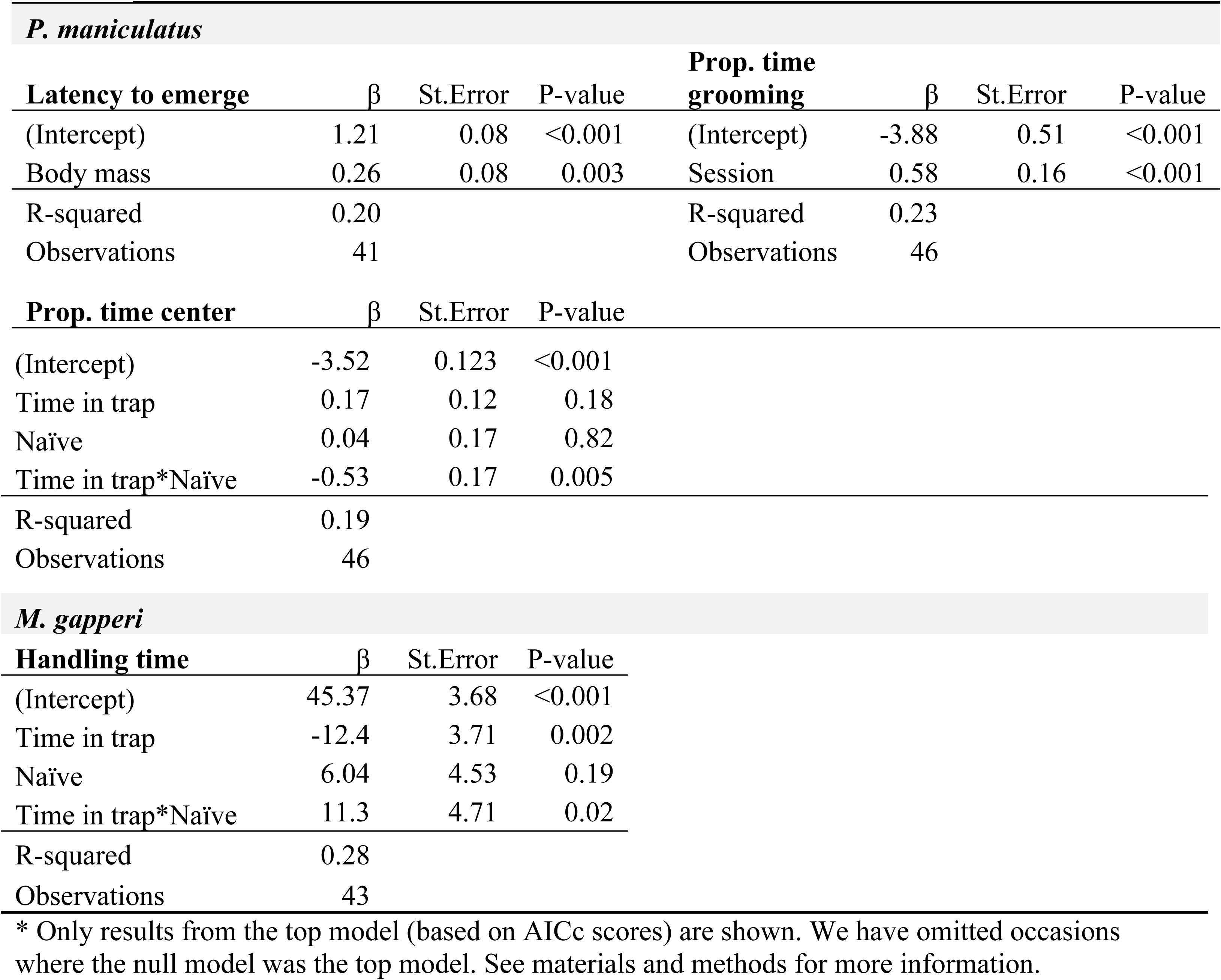
Model output of top-ranked linear models* predicting behaviors performed during standardized tests in deer mice (*Peromyscus maniculatus*) and southern red-backed voles (*Myodes gapperi*).

| *P. maniculatus* |  |  |  |  |  |  |  |
| --- | --- | --- | --- | --- | --- | --- | --- |
| Latency to emerge | $\beta$ | St.Error | P-value | Prop. time grooming | $\beta$ | St.Error | P-value |
| (Intercept) | 1.21 | 0.08 | <0.001 | (Intercept) | -3.88 | 0.51 | <0.001 |
| Body mass | 0.26 | 0.08 | 0.003 | Session | 0.58 | 0.16 | <0.001 |
| R-squared | 0.20 |  |  | R-squared | 0.23 |  |  |
| Observations | 41 |  |  | Observations | 46 |  |  |
| Prop. time center | $\beta$ | St.Error | P-value | | | | |

|  |  |  |  |
| --- | --- | --- | --- |
| (Intercept) | -3.52 | 0.123 | <0.001 |
| Time in trap | 0.17 | 0.12 | 0.18 |
| Naïve | 0.04 | 0.17 | 0.82 |
| Time in trap*Naïve | -0.53 | 0.17 | 0.005 |
| R-squared | 0.19 |  |  |
| Observations | 46 |  |  |
| ***M. gapperi*** |  |  |  |
| **Handling time** | **$\beta$** | **St.Error** | **P-value** |
| (Intercept) | 45.37 | 3.68 | <0.001 |
| Time in trap | -12.4 | 3.71 | 0.002 |
| Naïve | 6.04 | 4.53 | 0.19 |
| Time in trap*Naïve | 11.3 | 4.71 | 0.02 |
| R-squared | 0.28 |  |  |
| Observations | 43 |  |  |

\* Only results from the top model (based on AICc scores) are shown. We have omitted occasions where the null model was the top model. See materials and methods for more information.

**Fig 4.**
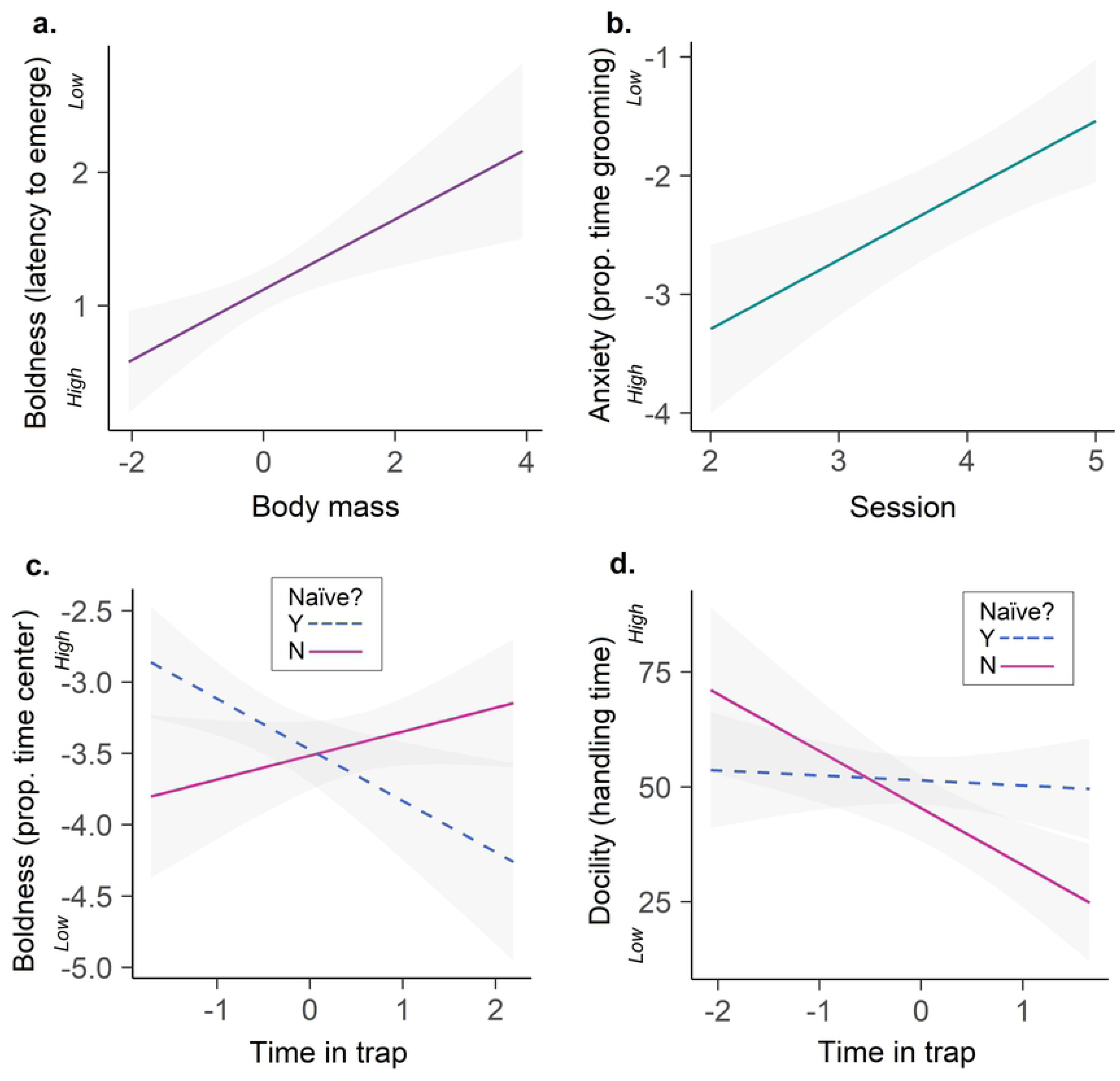
Factors predicting repeatable behaviors performed in the open-field test in deer mice (*Peromyscus maniculatus*) and southern red-backed voles (*Myodes gapperi*). **(a)** Deer mice with greater body mass took longer to emerge from the emergence test. **(b)** Trapping session influenced the proportion of time deer mice spent grooming in the open-field test (2 refers to July and 5 is October). **(c)** Deer mice who were naïve to trapping showed a negative relationship between time in the trap and the proportion of time spent in the center portion of the open-field test. Non-naïve mice showed the reverse relationship. **(d)** Voles who were not naïve to trapping showed a negative relationship between time in the trap and handling time. Results were obtained from linear models, and 95% CI from the models are shown. Variables “time in trap” and “body mass” have been z-standardized, and the variables “latency to emerge”, “prop. time grooming”, and “prop. time center” are on a log10 scale.

## Discussion

We studied the effects of live trapping on behaviors performed during three standard behavioral tests in deer mice and southern red-backed voles. Our major findings were that for these species, 12 out of 14 behaviors exhibited during routine behavioral tests were not affected by the amount of time that individuals had spent confined in traps. In the two instances where the time spent confined in traps did predict behavior, effect sizes were relatively small, and the direction of the relationship was different for individuals who were naïve to trapping than those who had been trapped previously, indicating that an individual’s previous experience with a trap interacts with this process. Overall, these results suggest that personality data collected from wild, trapped small mammals is not confounded by the trapping process and, where an effect might be present, the predictive power of the time spent confined to traps is relatively weak and possibly not affecting the overall interpretation of results.

Previous research has not explored the effects of live trapping on personality measurements, however, studies investigating the impacts of live trapping on hormonal stress responses have had mixed findings. Specifically, it has been shown in southern red-backed voles and meadow voles that live trapping induces an initial stress response, but that this response is not heightened following prolonged confinement inside traps (29,36). In our study, the observed behavior of red-backed voles in behavioral tests was consistent with these findings and 6 out of 7 behaviors showed no correlation with the time that the animal had spent previously confined inside of a trap. Previous studies investigating the correlation between stress response and duration of trap confinement in deer mice saw that after prolonged time spent in traps, stress hormone levels were significantly higher than after a short duration of trap confinement (36). By contrast, our results show no correlation between 6 out of 7 behavioral measurements and trap duration in the deer mouse. Although a hormonal change does not necessarily precede a change in behavior, we would expect to see an observable behavioral change in individual deer mice experiencing elevated glucocorticoid levels (for example, by affecting behaviors that indicate activity level such as speed of locomotion and rearing). Instead, the one behavior in deer mice for which “time in trap” occurred in the top model was the proportion of time spent in the center of the open-field test, a behavior which is most commonly interpreted as indicating the degree of boldness (Table 1). Interestingly, our results show that individuals who had never been trapped previously behaved more boldly in the open-field test (spending more time in the center portion) when their confinement duration was short rather than long. Individuals who had been trapped at least once previously showed the opposite effect; bolder behavior was seen in animals who had spent longer durations in the trap than those who had spent shorter durations (Figure 4c.). In voles, the one behavior that was affected by the “time in trap” was handling time, or the amount of time spent immobile during a one-minute handling bag test. This behavior is commonly used to assess docility (Table 1). Our results showed that for non-naïve individuals only (i.e., only those who had been trapped at least once previously), shorter durations in the trap correlated with increased docility (Figure 4d.).

Since 86% of observed behaviors by deer mice and voles showed no correlation with the variable “time in trap”, and all four variables indicating activity showed no correlations, we suspect that the duration of trap confinement is not providing a prolonged stressor for small mammals. It may be noteworthy that the previous trap response studies of deer mice and voles used Sherman traps instead of the Longworth traps used in this study. Longworth traps differ from Sherman traps in that they have a separate nest chamber (providing additional warmth and protection). Additionally, we took steps to limit stress by ensure that bedding remained dry (i.e., limiting trapping in adverse weather and replacing damp bedding immediately), and providing ample bait inside the traps. Further, we checked traps twice a day to limit confinement durations. We can’t speculate on whether these precautions were adequate in our study to stop a subsequent release of glucocorticoids after the initial stressor of the trapping event, but regardless, prolonged confinement in a Longworth trap does not seem to result in an observable change for the majority of behaviors in either study species. Future research examining this relationship in other species and other study populations will help to assess and confirm the generalizability of these findings. In the two cases where “time in trap” showed relatively weak predictive power, both arose as an interaction with the variable “naïve”. We suggest that other studies investigating personality in small mammals control in analyses for whether or not animals have been captured previously.

An animal’s personality depicts its unique way of experiencing the world and coping with life’s challenges (3). Using standardized behavioral tests, it is possible to capture different components of an individual’s complex personality, for example by observing activity levels and interactions with novel objects and environments (33). Our results show some evidence that an individual’s behavior in standard tests can be predicted in part by body mass and seasonality (Figure 4). Specifically, we found that heavier deer mice were slightly more timid than lighter mice (seen in their longer latencies to emerge from the emergence test), and that mice groomed more (indicating coping) in the autumn than they did in the early and mid-summer. These models showed low fit to the data; suggesting that the complexity of an individual’s personality is a difficult thing to predict.

Personality studies on wild populations will likely continue to become more common as further research demonstrates the cascade-effects that individual behavioral traits can have on populations and communities (14,16,18,19,68). Hence, it is critical to ensure that the very process we seek to illuminate is not being confounded by our methods of obtaining data. Our findings provide evidence that time spent inside of Longworth traps does not determine behaviors performed during standardized tests in two different small mammal species. Therefore, our results suggest that personality measurements on wild, trapped small mammals are not regulated by trapping procedures.

## Acknowledgments

This work was supported by the USDA-National Institute of Food and Agriculture McIntire-Stennis projects (ME041620 and ME041913) through the Maine Agricultural and Forest Experiment Station. We also thank a number of dedicated field and lab technicians for helping with data collection and video analysis, Bryn Evans for help with map preparation, Malcolm Hunter for comments on a previous version of this manuscript, Laura Kenefic (U.S. Forest Service), and Keith Kanoti for maintaining and facilitating research at the Penobscot Experimental Forest. A special thanks goes to Sara Boone for support in the field.

## Author contributions

AM, ST, and AMB conceived and designed the experiment. ST and AMB performed the experiment. AMB analyzed the data and wrote the first draft of the manuscript. All authors contributed to the final version of the manuscript.

